# Uniformity of spheroid-on-chip by surface treatment of PDMS microfluidic platforms

**DOI:** 10.1101/2022.02.15.480543

**Authors:** Neda Azizipour, Rahi Avazpour, Mohamad Sawan, Derek H. Rosenzweig, Abdellah Ajji

## Abstract

Spheroids have emerged as a more reliable model for drug screening when compared with 2D culture models. Microfluidic based biochips have many advantages over other 3D cell culture models for drug testing on spheroids, including precise control of the cellular microenvironment. The control of the cell adhesion to the surface is one of the most important challenges affecting the size and the geometry of the spheroids which could be controlled by appropriate surface engineering methods. We have studied the modification of the PDMS surface properties treated by applying different concentrations of the two anti-fouling coatings (BSA and Pluronic F-68). The desired treatment of PDMS surface effectively inhibits cell adhesion to the surface and promotes cells self-aggregations to form more uniform and healthy spheroids for a longer period of time. The microscopic observations with qualitative and quantitate data revealed that surface properties drastically affect the number of the spheroids formed on-chip and their geometry. We used human breast cancer cell line (MDA-MB-231-GFP) while the concentration of the chemical coatings and incubation time were adjusted. Proper repellent PDMS surfaces were provided with minimum cell attachment and facilitated spheroid formation when compared with non-treated PDMS. The results demonstrate fundamental and helpful patterns for microfluidic based cell culture applications to improve the quantity and quality of spheroid formation on-chip which are strongly manipulated by surface properties (i.e., morphology, roughness, wettability and etc.)

## 1. Introduction

The tumor microenvironment is known to play a crucial role in the development of cancer [1, 2]. It is therefore important to mimic the in vivo like tumor microenvironment in vitro to better study the mechanisms of cancer progression and enhance new therapeutics. Studies of the cancer mostly relay on 2D cell culture models using immortalized cell lines [1]. However it is known that signals from extracellular matrix and 3D tumor microenvironment play crucial role in the maintenance of tissue specifications [3]. Nevertheless, tumor cells respond to chemotherapy and environmental cues differently when they are cultured in 3D microenvironment [1, 4]. 3D cell culture has gained much traction over the past decades with its evident advantages of more physiologically relevant data and more predictive responses in drug discovery and disease modeling [5]. Among various 3D cell culture models, spheroids are cell colonies formed by self-assembly which act as excellent physiologically relevant models to provide more reliable therapeutic readouts [6, 7]. Spheroids can be made by cancer cell lines or cells isolated from patients [8, 9]. These *in vitro* models can reproduce some of the physiological aspects of *in vivo* tumors such as non-uniform distribution of nutrients and oxygen, limited availability of nutrients and/or drugs in their center, various metabolism and proliferation of cells in different part of the spheroids and being drug resistance [6]. They also have closer gene expression profiles to native tumors when compared with 2D models [10]. However, formation and maintenance of spheroids are the two main challenges to for broad-range use in cancer studies. Spontaneous spheroid formation occurs when cell-cell interactions dominate over cell-substrate interactions. Hanging-drop, spinner flask culture and NASA rotary cell culture systems are the most popular methods to form spheroids [1].

Controlling the size and geometry of spheroids can directly influence spheroid viability and drug uptake profile [11]. Accordingly, developing platforms to simply provide size-controlled spheroids are crucially important. Microfluidics which is the science of working with fluids in very small quantities (between 10^−9^ to 10^−18^ liters) in a set of micro-channels with dimensions in the range of tens to hundreds of micrometers, have recently been employed by various research groups worldwide to generate spheroid-based tumor-on-chip models [12-14]. However, quantity and quality of spheroid formation in these platforms and the ability to provide long-term monitoring of cellular activities are still among the remaining challenges.

Numerous studies have demonstrated that material surface properties (e.g. surface wettability, surface chemistry, surface charge and roughness) influence protein adsorption on the surface which consequently regulates cell adhesion, cellular communications and their proliferation [15]. Different materials have been used to fabricate microfluidic-based cell culture devices [16, 17]. In the past few years, Poly dimethylsiloxane (PDMS) has attracted more interest in microfluidic biochip fabrication due to its unique properties and particularly because of its biocompatibility, transparency, ease of fabrication and rapid prototyping [17]. However, cell culture results on the PDMS biochips greatly depend on the physiochemical properties of the PDMS surface [18]. PDMS is inherently hydrophobic due to a high surface energy barrier which may cause challenges for the flow of aqueous solutions into the channels [1]. It is mostly believed that hydrophobic surfaces have higher levels of protein adsorption than that of hydrophilic surfaces [1, 19]. Protein adsorption will promote cell adhesion on PDMS surface which consequently results in channel clogging in microfluidic based cell culture platforms [19, 20]. Various methods have been used by researchers to reduce cell attachment on the PDMS surfaces, yet full optimization remains to be determined.

Plasma oxygen is widely used in microfluidic devices to permanently bond PDMS layers to each other or into another surface (e.g. a glass slide). Plasma oxygen introduces hydroxyl groups on the PDMS surface temporarily to render the surface hydrophilic and increase in electro osmotic flow (EOF). This leads to facilitated fluid flow into the micro channels. However, the treated PDMS surfaces can undergo hydrophobic recovery by aging time due to the migration of low-molar-mass PDMS species from the bulk to the surface [17]. Alternatively, surface modification with non-ionic surfactants including poly (ethylene oxide) (PEO)-terminated triblock polymers (e.g. Pluronic) [21] and modification with blocking molecules with strong anti-fouling properties (e.g. bovine serum albumin (BSA)) [22] has been widely used to “block” protein adsorption and cell adhesion on the PDMS surface. Chemical surface modifications and coatings to reduce protein adsorption and cell attachment have been studied earlier elsewhere [21, 22]. However, the correlation of surface chemistry, surface morphology and surface wettability with cellular response to a surface, the 3D morphology of cell aggregations and their growth pattern remain unclear. Moreover, most of the previous studies did not employ the effectiveness of their techniques to prevent or to enhance cell adhesion to the surface on cell culture platforms or they only relay on 2D culture models. There are also contradictory results declared in the literature concerning cell tendency to hydrophobic or hydrophilic surfaces. It is therefore, of paramount importance to assess the effectiveness of surface treatments in the complex 3D environment for which they are intended.

To the best of our knowledge, no investigation has utilized the correlation between broad ranges of surface properties with cell responses to the surface in 3D model. Accordingly, a 3D microfluidic microenvironment for spheroid formation on-chip has been employed in this work. In this study, wetting properties of the PDMS surface, after surface treatment with the two commonly used chemical coatings which have strong antifouling properties has been investigated by using water contact angle (WCA) measurements. Atomic force microscopy (AFM) imaging was used to offer nano-scale insights on the surface morphology of the PDMS after treatment leading to surface properties changes. For the first time, we investigate and show the correlation between the concentration of the chemical coating and their incubation time to provide different degrees of surface wettability and morphology and their influence on cell adhesion to the surface and quality and quantity of the spheroid formation on biochip. Herein, we present a set of works to understand how physicochemical properties and the surface engineering of PDMS alter spheroid formation on biochip, to guide future on-chip cancer study. We anticipate that this method can be lead to efficient surface treatment in microfluidic-based biochips for cancer study.

## 2. Materials and methodology

### Fabrication of PDMS layer

Constant three-millimeter thick PDMS layer were cured by mixing elastomer base and curing agent with the ratio of 10:1 (w/w) (Sylgard 184 Silicone elastomer kit, Dow Corning). The PDMS mixture was poured in a petri dish. It was then degassed for 1 hour under vacuum to remove bubbles and cured at 65 °C in the oven for 2 h. When the PDMS samples cured, they were cut in approximately 2.5 cm × 2.5 cm pieces.

### Surface Modification of PDMS layer

Surface modification process has performed in two steps as described below:

#### Treatment of PDMS layers with oxygen plasma

The cured PDMS layers were exposed to oxygen plasma (liquid air, alphagaz 1, 99.999% purity) which was set at a flow rate of 20 sccm (standard cubic centimeters) controlled by a mass flow controller (MKS 247C channel readout MKS 1259B-00100RV (0-100 sccm). The pressure in the chamber (Cylindrical with a 9” diameter and 1.25” height) was set at 600 mTorr (80 Pa) and fixed using a butterfly valve during the process. The plasma was initiated by radio frequency plasma at 13.56 MHz (ENI model HF-300 impedance matching unit Plasma therm AMNS 3000-E) and the applied power was set at 20 Watts, with a plasma exposure time of 20 seconds per sample.

#### Surface treatment with chemical coating

To provide integrity of data, all the samples have been treated with chemical coating on the day 7^th^ after plasma exposure. Samples surfaces were washed gently with Ethanol 99% and right after samples were immerged in their chemical coating solutions. In this study, we have chosen two different coating with three different concentrations of each (6 different coatings in total). BSA and Pluronic F-68 which are commonly used surface coating to reduce cell adhesion to the surfaces have chosen for this purpose. Chemical coatings were prepared in concentrations of 3% w/v, 5% w/v and 10% w/v of each, to treat sample surfaces. Two time points have been used in this study. One set of samples have been treated by immersion in freshly prepared BSA solution and Pluronic F-68 solution for approximately one hour at 37 °C and another set of samples have been treated over night at 37 °C to ensure maximum adsorption.

### Surface characterisation

#### Water contact angle (WCA) measurement

After surface treatment, each sample was gently washed with HBSS (Wisent Bioproducts, St-Bruno, QC) to remove the excess BSA and Pluronic on top of the PDMS, also to access if the WCA changes and morphology changes on the PDMS surface were long term or just visible due to excess Pluronic and BSA crystals on the surface.

Right after surfaces have washed with complete cell culture media to follow the same procedure right before the cell culture on PDMS biochips.

WCA were measured directly on clean and flat surfaces with a goniometer (Data Physic OCA, SCA20 software) using the sessile drop technique. An intermediate equilibrium of water-air-solid contact angle was obtained by depositing a 2 μl water droplet on the surface using the calibrated syringe. A photograph of each droplet was taken 30 sec after contact with the surface by using the VCA Optima contact angle analyzer (AST products, Billerica, MA) to study the hydrophobicity. Water contact angle measurements were measured in triplicate for each sample on the contact side.

#### Scanning electron microscopy (SEM) analysis

Topography of the treated PDMS surface and non-treated PDMS were recorded on a field emission scanning electron microscope (JSM 7600F, JEOL) using acceleration potential of 15 kV. SEM images were captured in resolution of 1280 × 1240 pixels.

#### Atomic force microscopy (AFM) measurement

The topography and roughness of treated PDMS surfaces along with non-treated surface were studied using an Atomic force microscopy (AFM). All AFM images were captured in the air at room temperature using tapping mode on a (Dimension ICON AFM, Bruker/Santa Barbara, CA). Intermittent contact imaging (i.e., “tapping mode”) was performed at a scan rate of 0.8Hz using etched silicon cantilevers (ACTA from AppNano) with a resonance frequency around 300kHz, a spring constant of ≈ 42 N/m, and tip radius of ‘lt;10 nm. All Images were obtained with a medium tip oscillation damping (20-30%).

#### Assessment of the surface characteristics of PDMS

To repeat each experiment in minimum three replicates, numerous PDMS pieces have been fabricated with soft lithography and have been treated with oxygen plasma as described above. The hydrophobic recovery of PDMS after treatment with oxygen plasma occurs by aging time of plasma treatment due to the migration of lower molecular weight species to the surface [23]. Therefore, PDMS pieces have treated with chemical coating on the day 7^th^ after plasma treatment to provide same physiochemical properties for all the samples before coating with chemicals and to exclude the effect of plasma treatment on data interpretation after chemical modifications. Three groups of samples were provided for contact angel measurement, SEM imaging and AFM analysis. Each group was treated with three different concentrations (3%, 5% and 10%) of two chemical coatings: BSA and Pluronic F-68.

### Fabrication of microfluidic device to form spheroid on biochip

Each microfluidic device is composed of two PDMS layers obtained from 3D printed master mold. Design of the channels is adapted from Astolfi et al. [24] and the parameters of each channel and wells are broadly the same as the device described by Astolfi et al. [24], except the height of the wells have increased to 540 μm to optimize device operation [25]. The bottom layer of the device consists of 2 open channels with a 600 μm wide square cross-section. Each channel is containing of five 600 μm-wide square-bottom micro-wells of 540 μm in height. The top layer PDMS with 3mm diameter inlet and outlet holes bonded with Plasma Oxygen (same conditions described above for PDMS surface treatment with plasma oxygen) to the bottom layer. The polymeric resin molds (HTM 140 resin, EnvisionTEC GmbH, Gladbeck, Germany) have 3D printed using stereolithography printer (Freeform Pico and Pico 2 HD, Asiga, Alexandria, Australia).

Briefly, a mixture of PDMS (Sylgard 184 polydimethylsiloxane elastomer kit, Dow Corning, Midland, USA) base polymer to curing agent at a mass ration of 10:1 prepared and mixed well, degassed and poured into each resin mold and allowed to crosslink and cure for 2 hours in 65 °C oven. When it is cured, PDMS layer gently peeled off from the mold. Following surface treatment with oxygen plasma, the two layers of the PDMS have bond together to form closed microfluidic channels and assembled into the microfluidic device.

### Surface treatment of channels

Each microfluidic device was sterilized and air bubbles were removed from each channel by using 100% ethanol (Sigma-Aldrich). Then each device was treated with chemical coating in order to assess the role of surface coating to reduce cell adhesion into the channels and facilitate spheroid formation on-chip. As described above, two chemical coatings were used in this study for surface treatment. Triblock copolymer surfactant Pluronic F-68 (Sigma-Aldrich) and BSA (Sigma-Aldrich) which are the most common surface coatings in PDMS based microfluidic devices to prevent cell adhesion were prepared in three different concentrations of 3% w/v, 5% w/v and 10% w/v of each for surface treatment. We have divided the devices to two groups to be treated with the same surface coating for two different incubation times. One group of devices have been treated with each chemical coating for approximately one hour at 37 °C and the second group of devices have been treated over night at 37 °C. Simply, by using P1000 micropipette, each one of the chemical coatings has introduced through each channel inlet and washed the channel for three to four times. Then each channel was filled with the chemical coating for the specific time designed for each group (Less than one hour for group one and overnight for group two of devices) and placed in the incubator ((37 °C, 5% CO2, 95% ambient air).

### Cell culture and spheroid formation on-chip

MDA-MB-231-GFP cells (human mammary gland adenocarcinoma) were donated by laboratory of Professor M.Park (McGill University) and were maintained in high glucose Dulbecco’s Modified Eagle Media (DMEM, Sigma Aldrich) supplemented with 10% (v/v) Fetal Bovine Serum (FBS) (Sigma-Aldrich) and 1% (v/v) penicillin–streptomycin (Sigma-Aldrich) at 37 °C, 5% CO2 and 95% relative humidity (RH). Cells were seeded and passaged one or two times into 75 cm2 tissue culture flasks (Corning Inc., New York, USA) before spheroid formation on chip. 80–90% confluent cell cultures were used for the experiments on biochip. For spheroids culture on biochip, cells were washed with sterile phosphate buffered saline (PBS, Sigma Aldrich) and trypsinised with 0.25% Trypsin-EDTA solution (Sigma-Aldrich) to create a cell suspension. Right before the cell culture on chip. Then microfluidic devices removed from incubator and channels were rinsed three times with sterile HBSS (Wisent Bioproducts, St-Bruno, QC) to clean the channels from chemical coating residues. Then a cell suspension with the concentration of 5□×□10^5^ cells/mL was introduced into channels inlet by using P200 micropipette. Cell suspension will be flow into the channels using gravity-driven flow. The tube connected to the outlet will be lowered to approximately 10-15 cm below the inlet during cell seeding process. Accordingly, gravity resulted from this difference in height, will create a force to facilitate cell seeding. When the channels are completely filled with the cell mixture, flow was stopped and allowed cell sedimentation into the wells. The devices were then incubated under static conditions at humidified incubator with 5% CO_2_ at 37□°C for 24 hours. Spheroid formation was occurred by on-chip sedimentation and containment [12, 13]. During the cell loading process into the inlet of the channels, a portion of the cells will be trapped and settle down into the wells and the rest will flow into the rest of downstream wells or will be ejected through outlet. Since the surface of the channels were treated and became cell-resistant, cells self-aggregation to form spheroids were occurred inside each well within one day after cell seeding process. On first day post-seeding, non-adherent cells were washed off by gently rinsing the channels with complete media and cell aggregated spheroids remain inside the wells. Inlets and outlets of the devices filled with complete media and devices were kept in the incubator under static conditions for a period of 7 days for farther experiments with daily media exchange by using P200 micropipette.

### On-chip observation of spheroid formation and proliferation tracking

Spheroid formation and proliferation were imaged directly through the thin PDMS layer by using Epifluorescence inverted microscope (Axio Observer.Z1, Zeiss, Oberkochen, Germany) and sCMOS camera (LaVision, Göttingen, Germany) and the objective lens EC Plan-Neofluar 5x / 0.15 for duration of 7 days incubation. The size of the spheroids was determined by measuring their diameters after they were imaged by fluorescent microscopy.

### Image J and data analysis

As explained above, spheroid images were recorded by using Epifluorescence inverted microscope (Axio Observer.Z1, Zeiss, Oberkochen, Germany) and sCMOS camera (LaVision, Göttingen, Germany) with the objective lens EC Plan-Neofluar 5x / 0.15. To analyse the fluorescent images of spheroids’s size, ImageJ (National Institute of Health, Maryland, USA) was used. The brightness level of green fluorescent has obtained by a digit value from zero to maximum 255. The analysing area of the spheroid cell culture has been carefully selected on their 2D image. All images have high resolution more than 1900 dpi. The average mean diameter can be written as Dm= (Dmax+Dmin)/2 [26]. The average size of each spheroid was reported as mean diameter ± standard deviation. The circularity of the spheroid 3D cell culture has been defined by the Circularity=Dmin/Dmax around a single sphere (**Figure 9** Appendix). Brightness level ratio is equal to BLR=100*Brightness level/255. Fluorescent signals as the function of cell density in the spheroids were used for calculations. All Data have been reported as the mean□±□standard error (SE) of minimum three independent tests. All error bars in figures indicate SE. Experiments were repeated at least three times with duplicate culture microfluidic channels for each condition. One representative experiment is presented where the same trends were seen in multiple trials.

## 3. Results and discussion

Previous studies have demonstrated that surface treatments with BSA and poly (ethylene oxide)-poly (propylene oxide)-poly (ethylene oxide) (PEO/PPO/PEO) co-polymers (Pluronics) strongly have anti-fouling characteristics[27, 28]. Liu et el. [27] was showed that PEO/PPO/PEO co-polymers can modify the surface to completely prevent cell attachment. But in the present study, we have observed that the success in the surface treatment highly depends on the concentration of the chemical coating and their incubation time with the surface.

### Water contact angle

Contact angle measurements were measured for non-treated and treated PDMS surface with BSA and Pluronic F-68 coatings. As explained above, all chemical coatings were carried out on day 7^th^ after plasma treatment to have the integrity in data as the contact angels decrease immediately after the plasma treatment due to the presence of polar groups (e.g. SiO2, Si–OH and Si–CH2OH) on the surface [29].

It is observed that for both surface treatments (BSA and Pluronic F-68), WCA decrease by increasing the incubation time and increase in the concentration of chemical coatings. The results showed that WCA on BSA coated surfaces decreased to close to 60 degree, while Pluronic F-68 coated surface WCA decreased to around 82 degree.

We observed that WCA of PDMS surface covered with BSA and Pluronic F-68 is lower than that of the bare PDMS surface, which means that both of these surface coating materials reduce the hydrophobicity of PDMS.

Figure 1 demonstrates the WCA of PDMS layers in the contact side vs different concentrations of chemical coatings: A) The difference between water contact angles of PDMS coated after 1 hour and 24 hours of incubation with three concentrations of BSA, B) The difference between water contact angles of PDMS coated after 1 hour and 24 hours of incubation with three concentrations of Pluronic F-68. Error bars represent SE for 3 independent sets of experiments. Within each experiment, WCA was calculated by averaging data points on the same sample (n = 3).

**Figure 1.**
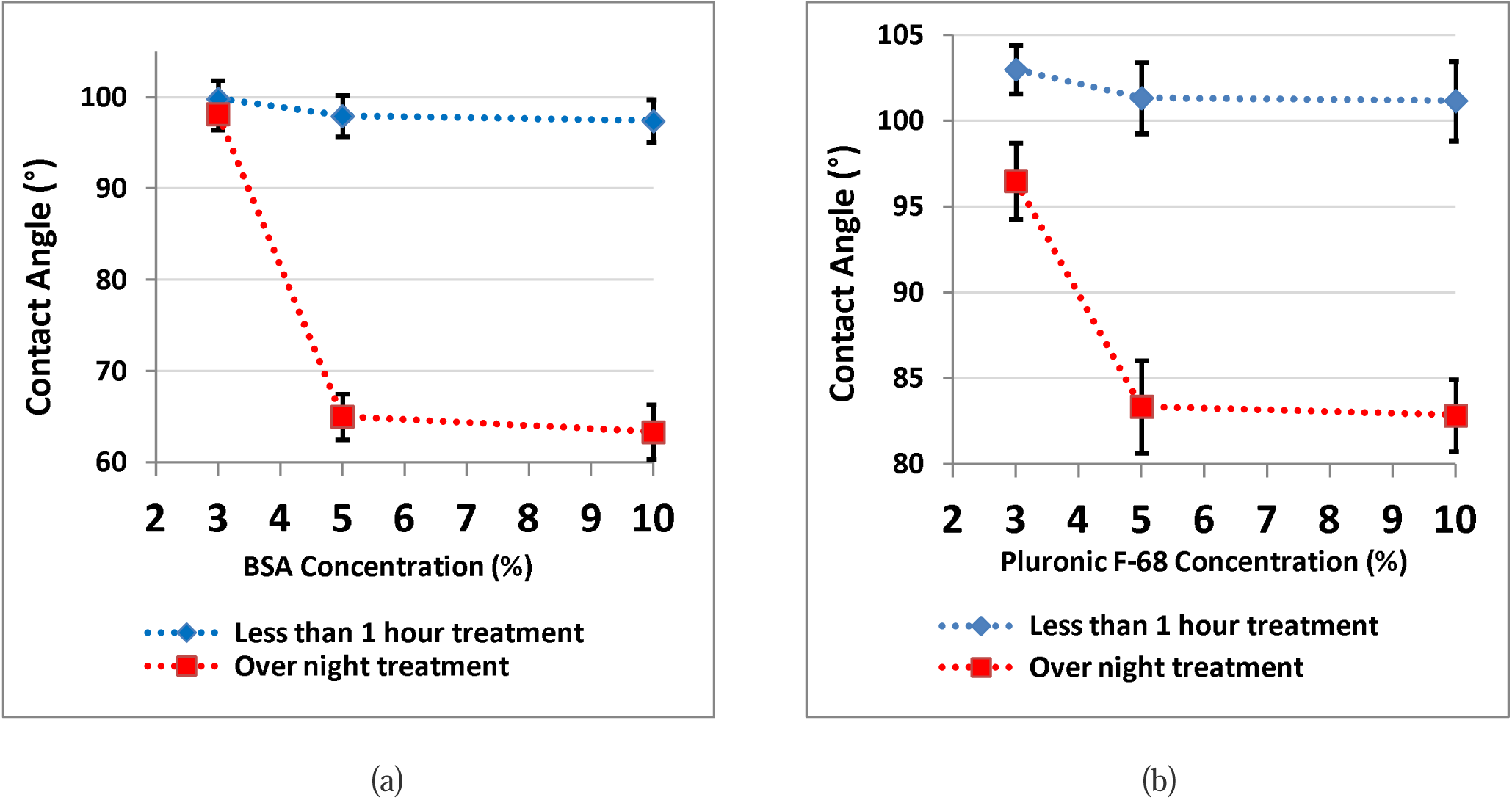
Measurement of WCA on the treated PDMS surface: (a) with BSA, (b) with Pluronic F-68

As decrease in WCA has been reported to reduce the protein adsorption to the surface, the results of this study suggest that PDMS surface treatment with both BSA and Pluronic F-68 that turned the chemical nature of the PDMS surface more hydrophilic, reduce cell attachment to the surface.

### SEM analysis

The results from SEM micrographs show that nano-structuring of PDMS have changed by surface treatment. Figure 2 Demonstrates results from surface treatment of PDMS with 5% Pluronic F-68 as an example to show the PDMS surface changes by surface treatment with chemical coating.

**Figure 2.**
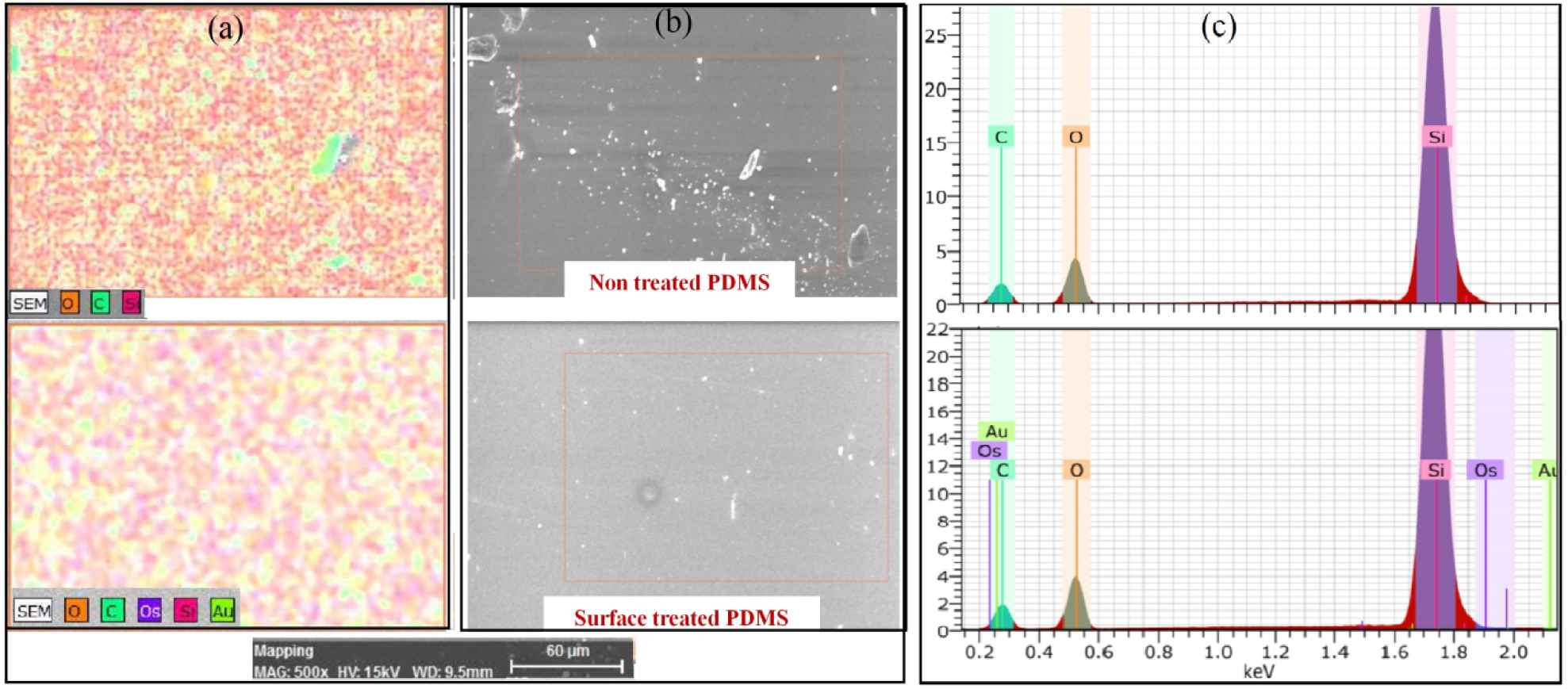
SEM of the non-treated and treated PDMS surface: (a) Modal color micrograph image, (b) SEM images, (c) Elemental peaks cps/eV [KeV]

The appearance of (C–O–C) peak and a hydroxyl peak (C–OH) for surface treatment with Pluronic F-68 and the presence of amide peaks on the PDMS surfaces treated with BSA (data is not presented in this work) indicate that the Pluronic F-68 and BSA molecules are adsorbed at the PDMS surface and modified the structure and morphology of the PDMS surface.

### AFM analysis

Regarding the capacity of different surface coatings for preventing unwanted cell adhesion to the surface, the surface coverage and the surface roughness are thought to be of the most importance.

In the present study, AFM was used to conduct a survey of the surface morphology of the three different concentrations of BSA and Pluronic F-68 as surface coating materials that are commonly used in the microfluidic based devices. The homogeneity and roughness of the surface coating was considered as an important parameter for this type of devices as it would impact the efficiency of cell repulsion.

When comparing the PDMS layers treated with different chemical coating in two different incubation times, a clear change in the surface morphology can be observed. For AFM image analysis to access the role of incubation time, we have chosen 5% concentration for each of BSA and Pluronic F-68. Incubation time designed as less than one hour and overnight treatment for each surface treatment.

AFM images of the non-coated and chemical coated PDMS surfaces have plotted in Figure 3.

**Figure 3.**
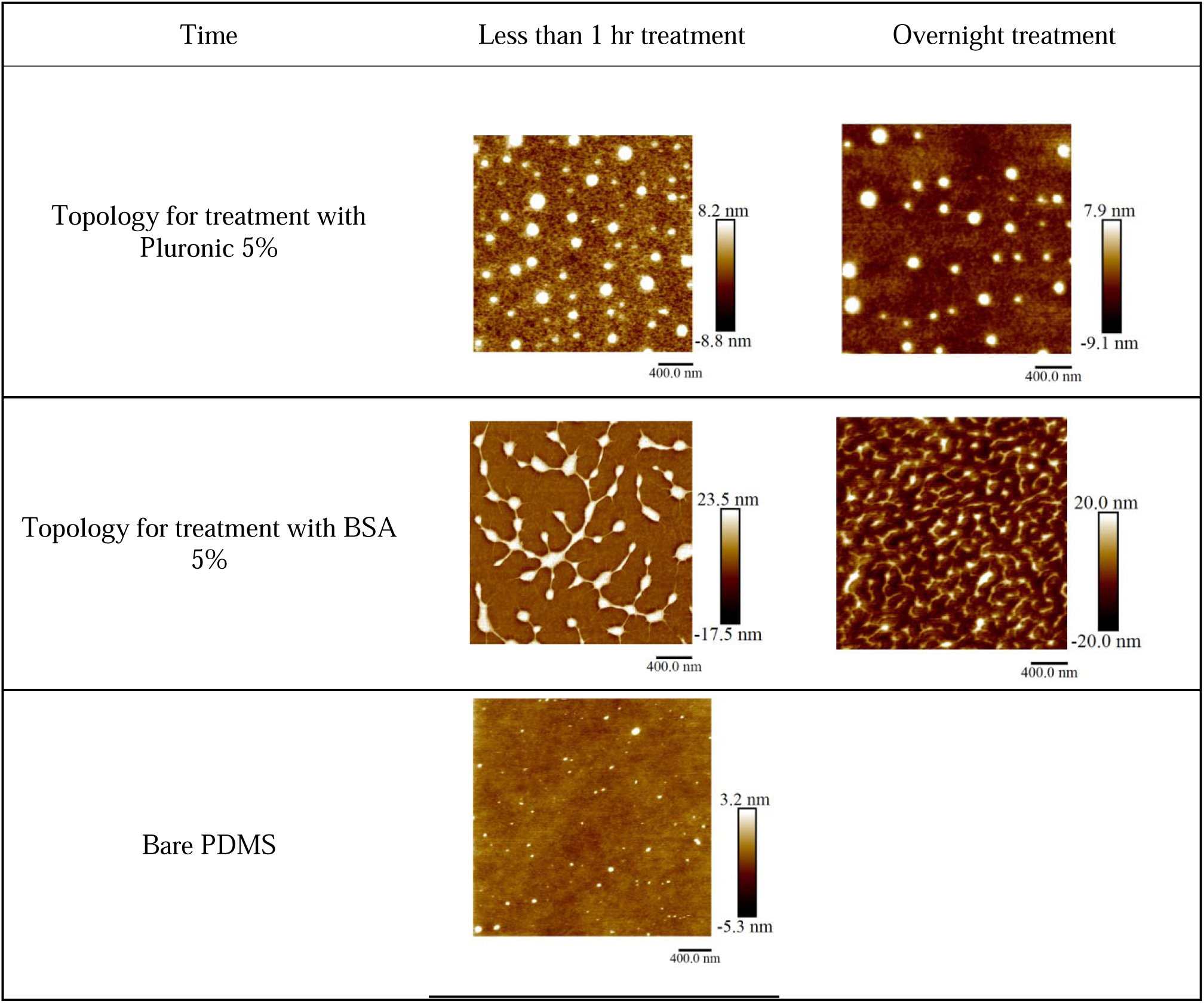
AFM images of the chemical coated and non-coated PDMS surfaces, for less than one hour and overnight treatment with BSA 5% and Pluronic F-68 5%, vs bare PDMS

As we expected, AFM images show the PDMS surface which is not treated, is homogenous and average roughness (Ra) is low. But the PDMS surfaces that have been modified demonstrated an increase in the surface roughness. PDMS surface covered with BSA has demonstrated a “peak and valley” structure which is probably due to protein unfolding upon the BSA adsorption. Then the internal hydrophobic domains can bind to other BSA molecules. Same pattern have observed by covering the surface with Pluronic F-68 that have increased the surface roughness.

In another set of experiments, PDMS surfaces have treated with three different concentrations of BSA and Pluronic F-68 overnight for AFM analysis. The results demonstrated that increasing the concentration, decrease surface roughness and increase the surface coverage. By increase in concentration, more molecules are available to cover the PDMS surface and accordingly they can fill out more the free spaces between peek and valley structures which will result in decrease in height differences between peek and valley structures and decrease in average topology values on the PDMS surfaces (Figure 4 and Table 2).

**Table 1.** Image J analysis of average surface topology value for short term and long term treatment with chemical coatings.

| Material | Short treatment with 5% agent | Long treatment with 5% agent |
| --- | --- | --- |
| Average topology (nm) for Pluronic | 8.3 | 7.9 |
| Average topology (nm) for BSA | 19.1 | 18.3 |

**Table 2.**
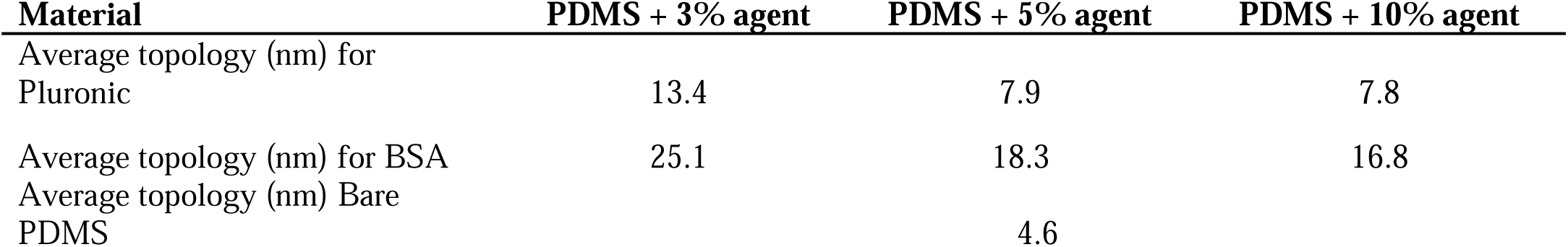
Average of topology value in nm for three concentrations of chemical coatings

| Material | PDMS + 3% agent | PDMS + 5% agent | PDMS + 10% agent |
| --- | --- | --- | --- |
| Average topology (nm) for Pluronic | 13.4 | 7.9 | 7.8 |
| Average topology (nm) for BSA | 25.1 | 18.3 | 16.8 |
| Average topology (nm) Bare PDMS |  | 4.6 |  |

**Figure 4.**
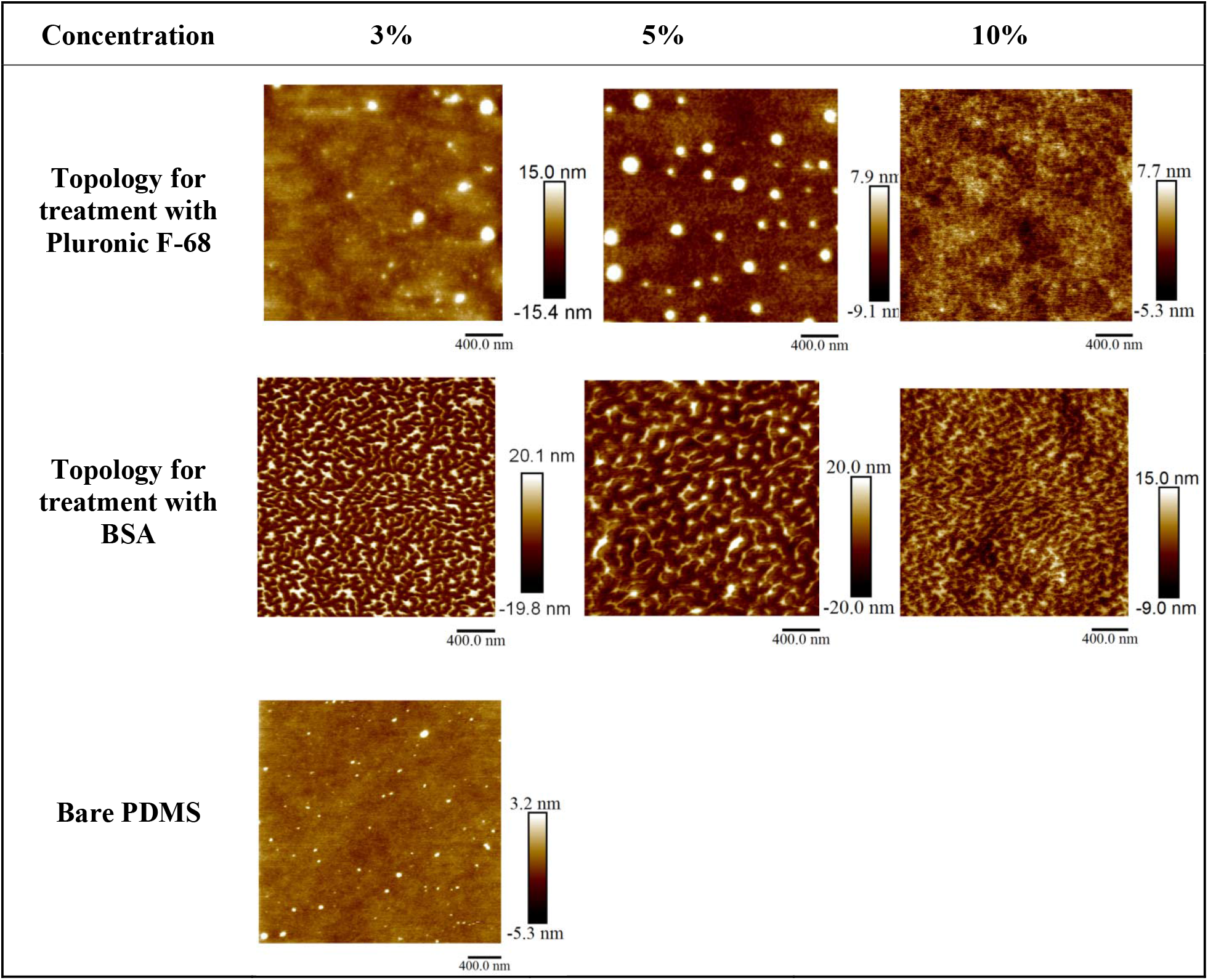
AFM images of the three concentrations of chemical coated PDMS surfaces with BSA and Pluronic F-68 vs bare PDMS.

### ImageJ analysis of treated PDMS

By quantifying the fluorescent intensity of the imaged surfaces using ImageJ software, the amount of adsorbed BSA or Pluronic on the PDMS surface was estimated. **Error! Reference source not found**. shows the results obtained by Image J.

Average of topology value in nm for three concentrations of chemical coating (BSA and Pluronic F-68) have calculated and summarized in Table 2. As observed, by increasing in the concentration of the chemical agents, average topology value, obtained by AFM analysis, decrease.

AFM images have analysed by Image J to evaluate surface coverage by each coating material.

For that purpose, we have measured the average of color intensity in AFM images produced by red, green and blue colors. Image analysis demonstrated that when the time of the surface treatment and concentration of the coating materials increase, the average number of red, green and blue color decrease. Accordingly, we interpreted that when the number is bigger, more peak structures are available in the surface and there are higher differences in height between peak and valley structures and surface covered with less material. However, when the number decreases, we interpreted that we have more surface coverage and less differences between peak and valley structures. Table 3 summarized the analysis by Image J.

**Table 3.**
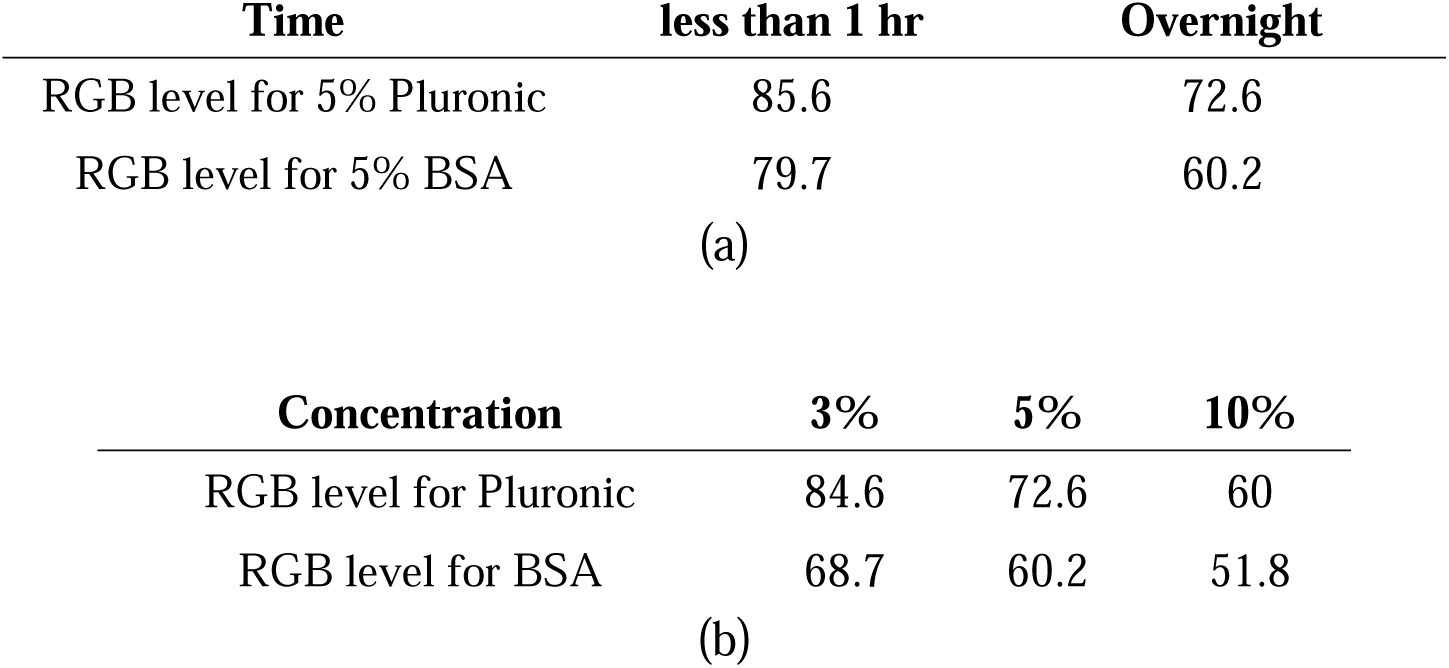
the results of AFM’s images analyzed by Image J, (a) the effect of treatment durations (b) the effect of concentrations

Table 3 summarizes the results of AFM’s images analyzed by Image J. RGB level is a number calculated by Image J and show the color intensity obtained from AFM images. R, G and B are representing red, green and blue color respectively. Decrease in RGB level number, demonstrates that we have more surface coverage and less height differences between peak and valley structures.

### Spheroid formation on microfluidic chip

This device is inspired by design developed by Astolfi et al. [24] to study MDTs. We have modified the design to have two separate microfluidic channels in each device. The height of each well is also modified according to the theoretical simulation [25] for optimal cell trapping. The cell mixture of MDA-MB-231 cells is introduced into the inlet of the each channel. Since the channels surface were treated and became resistant to cell attachment, a portion of the cells settled down into the wells by sedimentation and self-aggregated into cellular mass by secretion of their own local matrix and cell-cell contacts to form spheroids within one day in culture.

Each channel has five cell trapping micro wells where the cells trap by sedimentation and form the spheroids. The sizes of the spheroids were relatively uniform in each group and it mostly depends on the initial cell concentration of the cell mixture. We have experimentally optimized cell concentration to 5□×□10^5^ cells/mL in cell mixture for our device to obtain uniform size spheroids for duration of at least 7 days (Figure 5).

**Figure 5.**
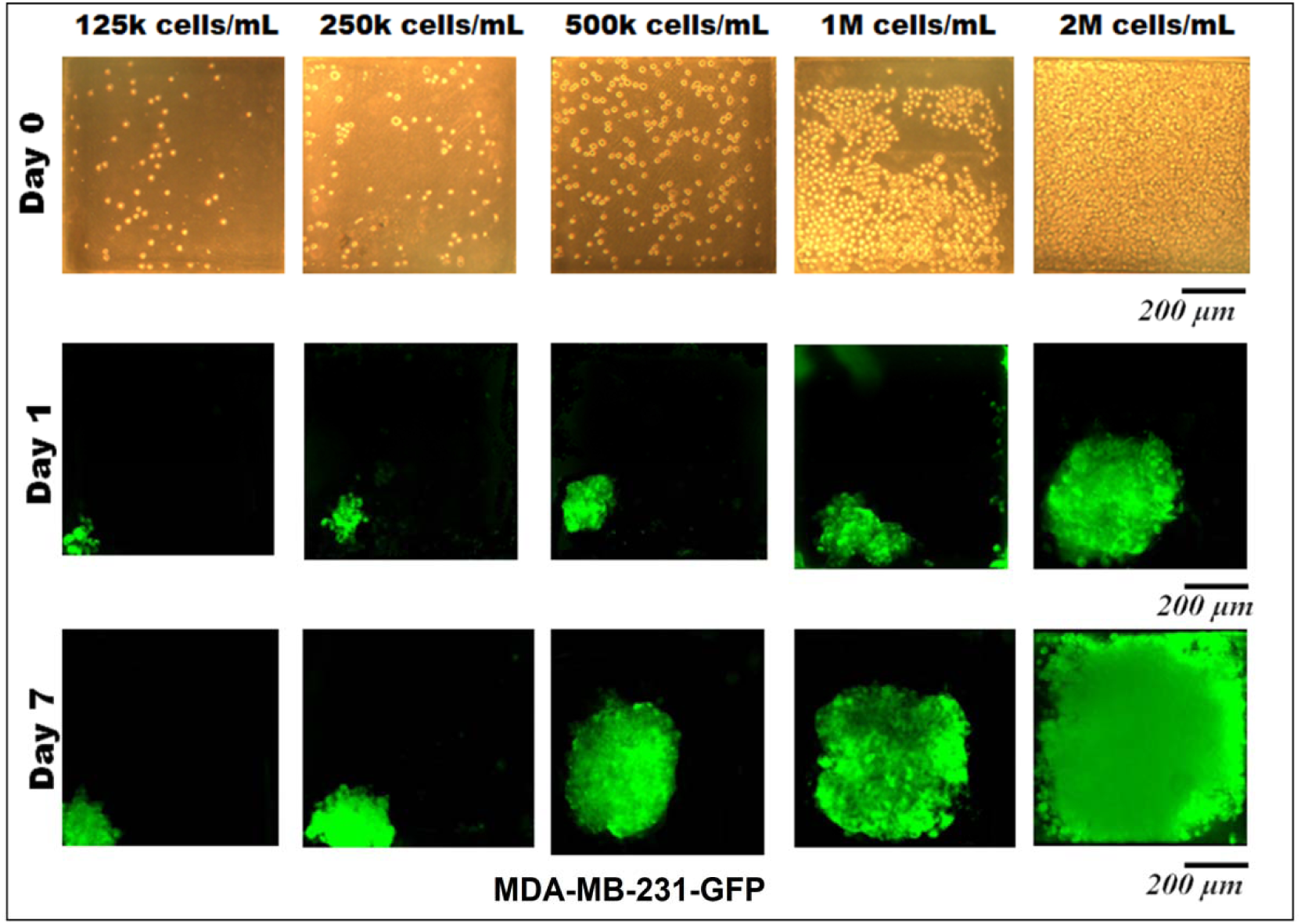
Spheroid formation on-chip with different initial cells concentration during 7 days. Fluorescent images of MDA-MB-231-GFP spheroid formation within microchannel. Different initial cells concentration applied to adjust the optimal initial cells concentration. (Size of chambers: 600, 600, 540 μm, Size of microchannel: 600 μm width, 600 μm height. Scale bar is 200 mm

**Figure 6.**
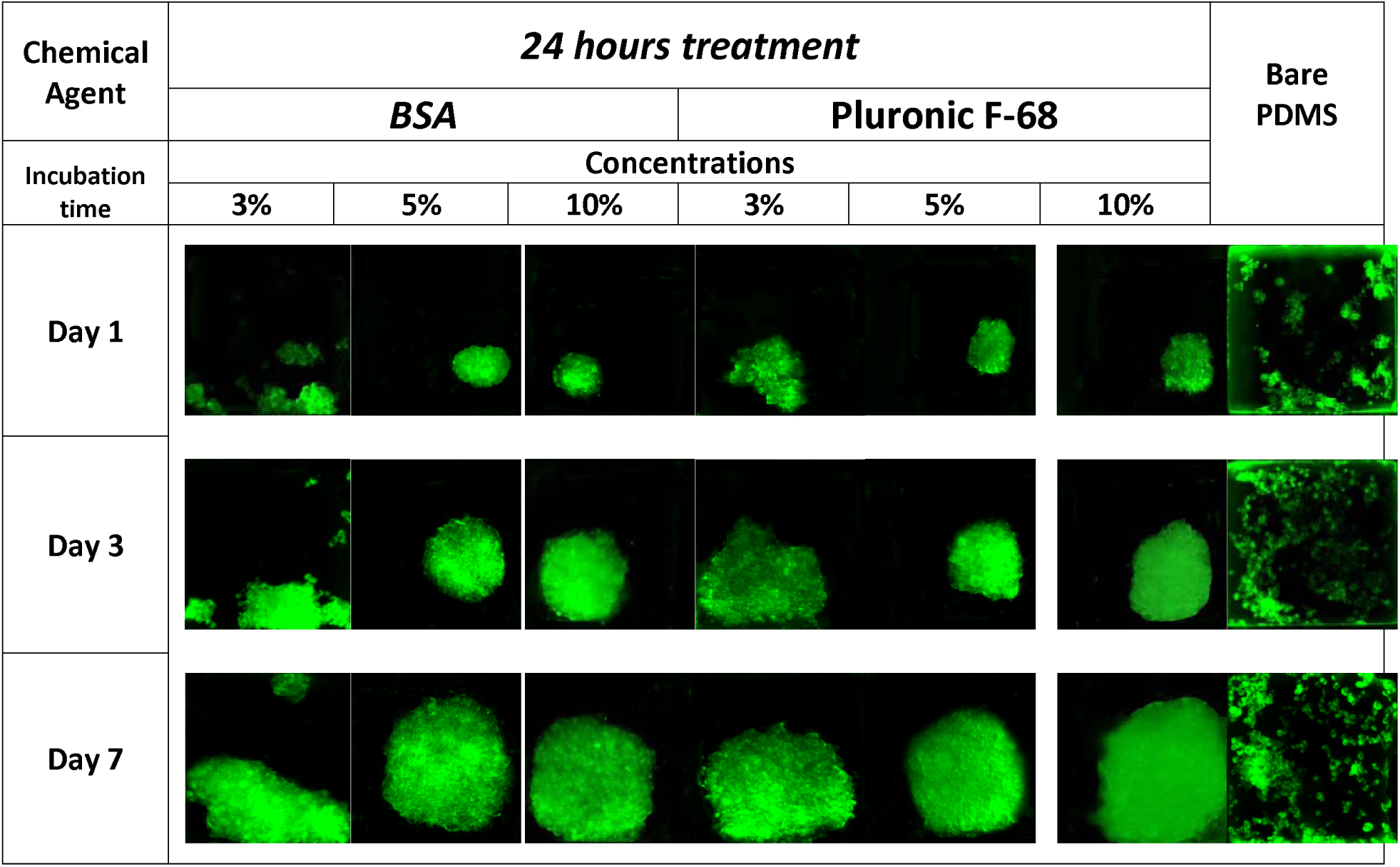
MDA-MB-231-GFP Spheroid formation on-chip treated with various concentrations of chemical coatings (BSA and Pluronic F-68) vs non-treated PDMS biochip. Images are representative of three independent experiments. Each square in this image is representing 600 μm*600 μm

In this device, the distance between each well provides sufficient amount of nutrients in a non-perfused system for one day according to COMSOL simulations (data are not provided).

Microscopic observations shows that there is no significant difference between the spheroids formed in different micro-wells through the channel and confirmed the uniform distribution of cells along the length of each channel. Fluorescent images of spheroids show the progressive development of spheroid sizes through 7 days of culture. Cells are capable to self-aggregate and form cellular clusters after one day of culture and transform into compact uniform-sized spheroids after day second of culture. **Error! Reference source not found**. demonstrates the cell aggregation process on non-treated biochip vs devices which have been treated with chemical coating (BSA and Pluronic F-68) to be resistant to cell adhesion into the PDMS surface. In contrast with biochips which have been treated overnight with chemical coatings, devices without any surface modifications and devices which only have been treated for a maximum of 1 hours of incubation with chemical coating were favourable for the attachment of MDA-MB-231 cells. In these devices cells tended to attach to the walls of the microfluidic channels and exhibited spread morphology which was propagated to adherent cell clusters attached to the surfaces of the micro-wells.

The spheroids morpholog**y** on the BSA-coated and Pluronic F-68-coated devices was observed to be very similar and it mostly depends on the concentration of the chemical coating.

As a control, cells were seeded on a PDMS device without any surface treatment. A significantly greater number of cell adhesions were observed on non-treated PMDS channels as demonstrated in the **Error! Reference source not found**. which illustrate the importance of chemical coating in prevention of cell attachment to surfaces.

Spheroids formed in surface treated PDMS biochips vs non-treated device. Cells tend to attach to PDMS surfaces and do not form spheroids inside the device. However, when the surface is treated with BSA or Pluronic F-68, surface become cell repellent and this promotes cells self-agglomerations to form spheroids. The growth pattern and viability of the MDA-MB-231 cells within the 3D environment over the course of 1 week is shown in this figure.

### Long term survival of spheroids on-chip

The percentage of spheroid formation on various surface treated biochips on day one and the survival rate of spheroids for a total of 7 days were tracked by fluorescence imaging. Based on data, the spheroids with average viabilities above 70% remain highly viable during the 7 days experimental period. In average 20% of spheroids remain viable between day 7 to day 14 (data not provided). However, spheroids size after day 7th demonstrated that MDA-MB-231 cells mainly remained quiescent and proliferated at a relatively slow rate. We suspect that the shape of spheroids contribute to their long viability. Spheroids with more spherical morphology stay highly viable for longer period of time probably due to smaller size of necrotic core in their center.

Spheroid formation by sedimentation on chemically coated PDMS chip, as described here is a simple and cost effective 3D cell culture method. However, cell clusters that rely on cells self-aggregation on devices with short term incubation with chemical coating are faster than long term (over-night) surface treatment with chemicals, but yielded cell clusters with non-uniform aggregations such as lobular structures with poor cell-cell contacts.

*Figure **7*** shows the growth pattern of the spheroids over the period of 7 days. We have used the Image J for analysing the growth pattern of the spheroids over the period of 7 days. The average values of three points from repeated tests have taken into the calculations. The brightness level of the green fluorescent color obtained by a digit value from zero to maximum 255 which respectively corresponds to darkest and brightest pixels. The analysing area of the spheroids have carefully selected on their 2D pictures. All the selected pictures have a high resolution more than 1900 dpi. The average mean diameter can be written as Dm=(Dmax+Dmin)/2 and circularity of the spheroids has been defined by the Circularity=Dmin/Dmax around a single sphere. Brightness level ratio is equal to BLR=100*Brightness level/255 in our calculations. The higher brightness levels correspond to higher uniformity and cell concentrations in the spheres.

**Figure 7.**
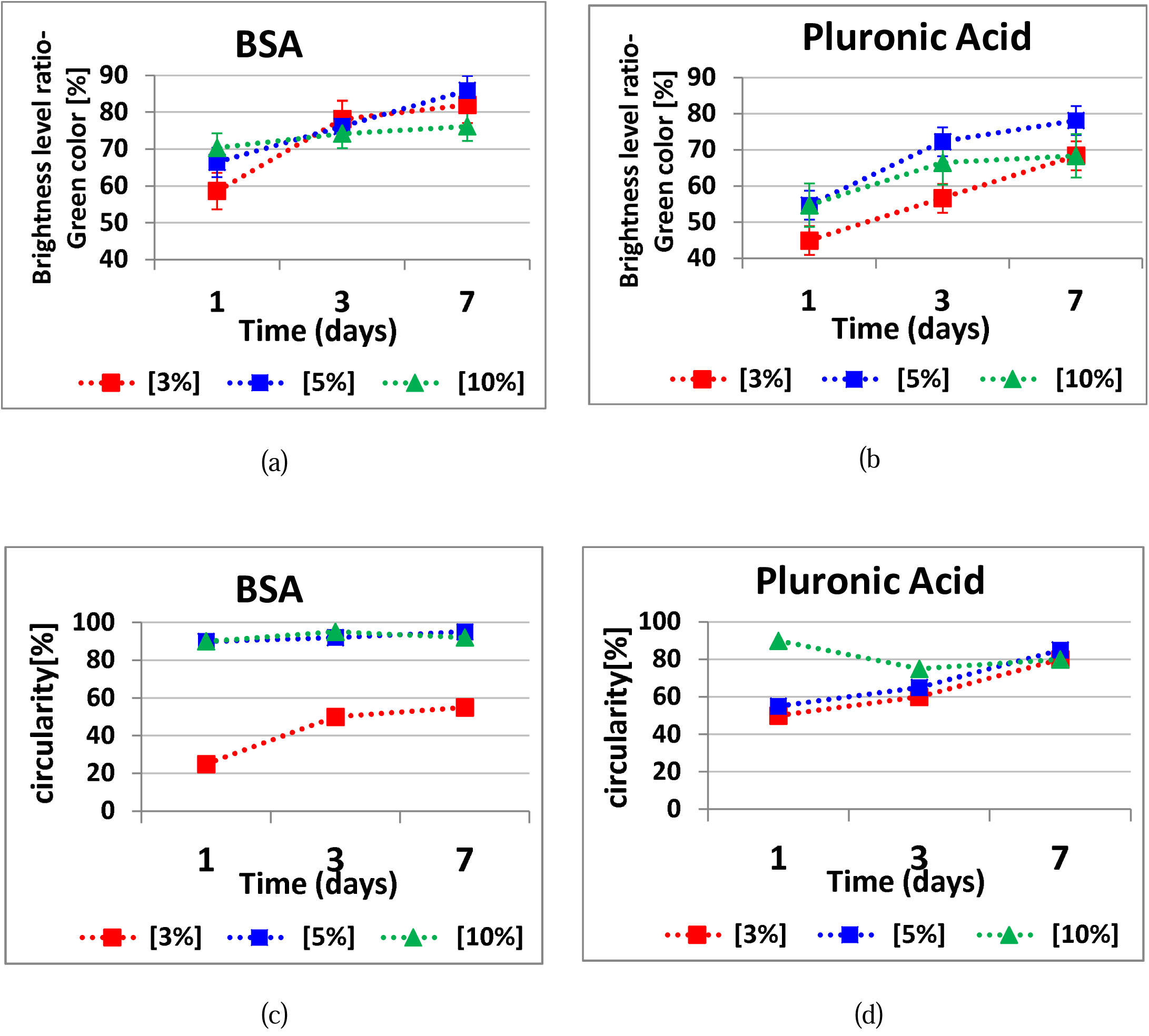
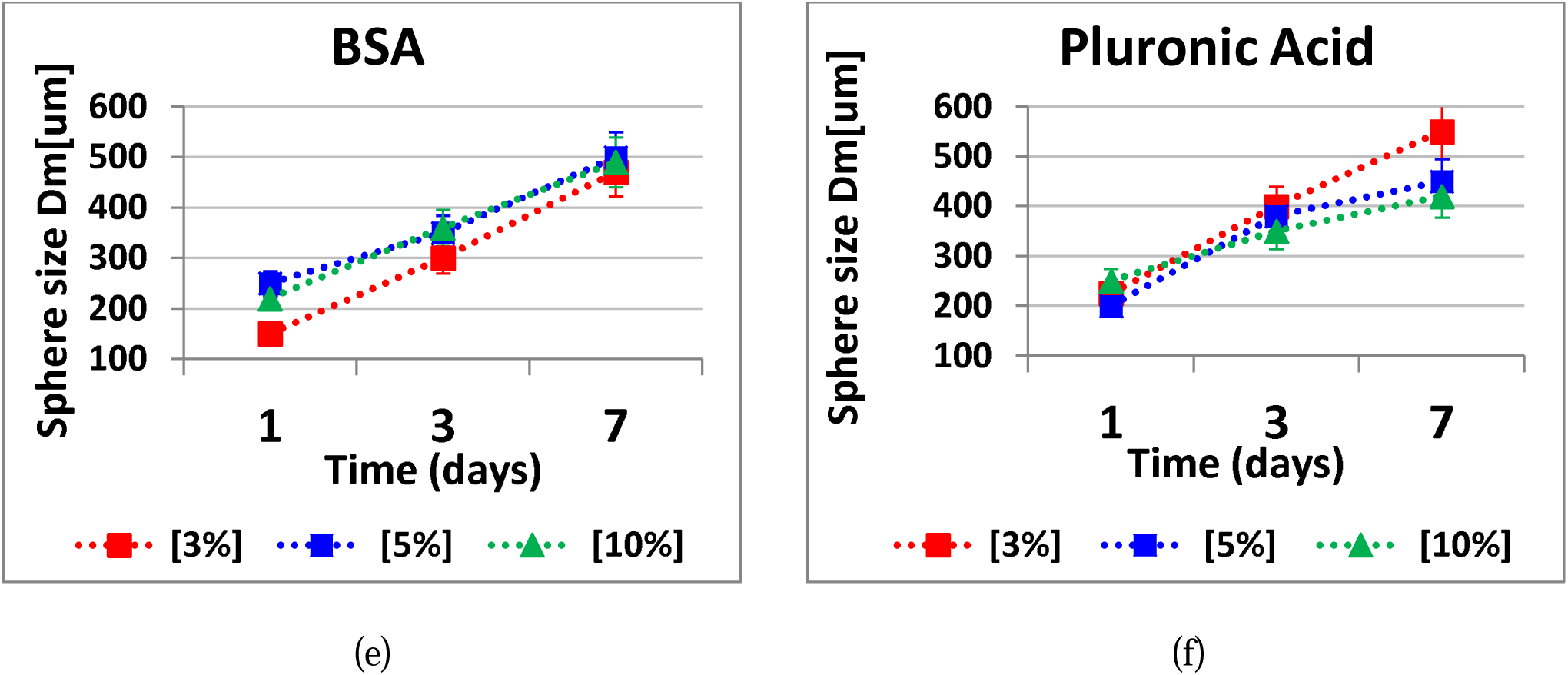
Spheroid analysis formed in PDMS biochip devices treated with different concentrations of BSA and Pluronic F-68 overnight Growth pattern of the spheroids analysed by Image J. a) and b) show the brightness level ration of green fluorescent for BSA and Pluronic F-68 respectively. c) and d) show the circularity of the spheroids which has defined as circularity as explained above for BSA and Pluronic F-68 respectively. e) and f) show the spheroid size in um. (Error bars represent ± SE, n=3).

Our data reveal that the BSA coating produced a pattern of cell repellent similar to that observed following Pluronic F-68 surface treatment. However, BSA was more effective starting at the concentration of 5% and the best results have recorded at the concentration of 10% to reduce cell adhesion to the surface. On the other hand, Pluronic F-68 has showed to be effective in reducing cell adhesion to surface starting at the concentration of 3%. According to our observation we believe that 5% is the optimal concentration for Pluronic F-68 to achieve higher percentage of uniformly sized spheroid formation on-chip as it seems that with 5% of Pluronic F-68, we are going to have enough surface coverage to prevent cell attachment to surface.

In our experiments, spheroid formation on surface treated with BSA 10% and 5% Pluronic F-68 was greater than to the other surfaces, suggesting that the moderate wettability (62-88 degree water contact angle) shared by these two surface coatings promote higher cell repellent properties for PDMS.

We believe that the concentration of the chemical coating and the incubation time are the two important factors to regulate surface chemistry and morphology of PDMS and provide the anti-fouling property for the surface. As in this study, we have focused on the chemical properties of the PDMS treated surfaces and the effect of other parameters was excluded. Accordingly, all the surfaces had the highest homogeneity as possible in terms of surface stiffness and etc. However, we know that the relation between surface properties and cell adhesion is complicated and it is not completely understood yet.

## 4. Conclusion

Understanding the correlation between surface properties and cellular responses to the surface is essential to design optimal surfaces in biomedical applications including microfluidic based biochip platforms. Although enormous studies have been done on PDMS surface modification, a systematic study on the simultaneous effect of anti-fouling coating on the PDMS surface properties and its effects on 3D cell culture on PDMS microfluidic chip platforms is still missing in the literature. As the cells behaviour is deeply influenced by the cellular microenvironment and the substrate they grow on, the objective of this study was to gain insight into the correlation among surface properties of PDMS based microfluidic platforms, cell responses to the surface and spheroid formation on-chip.

In this work, for the first time, PDMS surface modification for stably deterring cell adhesion to the surface and facilitate spheroid formation on-chip was compared quantitatively and qualitatively by using of two common anti-fouling coating materials: BSA as a traditional method and Pluronic F-68 as a PEO-based surfactant. The spheroid formation of MDA-MB-231 cells on PDMS microfluidic biochip following surface treatment with six different surface chemistries providing various degrees of surface wettability and morphology was evaluated.

This study provides not only fundamental understanding about control of cell behaviour in PDMS-based microfluidic platforms, but also a simple and cost effective method to effectively enhance the quality and quantity of spheroid formation on-chip for in vitro tests.

Our observations revealed that however surface treatment with BSA has resulted in different surface wettability and morphology when compared with surfaces treated with Pluronic F-68, the final spheroid formation on-chip resulted in a similar pattern following using both chemical coating modifications. Our data presented in this work demonstrated that moderately hydrophilic surfaces (62-88 degree water contact angle) promoted the highest level of cell repellent properties. In this study, we have produced greatly viable and uniform structure spheroids with the highest number of spheroid formation on-chip. We have focused on breast cancer cells in this work, although the technology described here is versatile and can be used for other cell types in their more physiologically relevant state. On the whole, our results confirm the importance of the surface modification of microfluidic based platforms to easily block non-specific cell adhesion to the surface and enhance the quantity and quality of spheroids on-chip for drug development and cancer studies.

In summary we have shown a simple, effective, practicable and repeatable method to modify the PDMS surface of biochips that can prevent cell adhesion into the surface and promote spheroid formation on biochips without the need of expensive additional materials (e.g. hydrogel, matrigel and etc.) to the cells to form spheroids on biochip for drug testing.

## Conflicts of Interest

The authors declare no conflict of interest.

## Acknowledgements

The authors gratefully acknowledge funding support from the Natural Sciences and Engineering Research Council (NSERC) of Canada discovery grants. The authors also gratefully acknowledge Professor Thomas Gervais (Polytechnique Montreal) for fruitful scientific discussions to design and fabrication of the biochip, and Professor Morag Park (McGill University) for kindly providing the cell line used in this study. N.A gratefully acknowledges the scholarship from Fondation et Alumni de Polytechnique Montréal donated by Royal Bank of Canada (RBC).

## List of Acronyms

AFM: Atomic force microscopy
BSA: Bovine serum albumin
EOF: Electro osmotic flow
HBSS: Hank’s balanced salt solution
PBS: Phosphate buffered saline
PDMS: Poly dimethylsiloxane
PEO: Poly (ethylene oxide)
Ra: Average roughness
SD: Standard deviation
SE: Standard error
SLA: Stereolithography apparatus
3D: 3 dimensions
2D: 2 dimensions
WCA: Water contact angle

## Appendix

BSA which is a heart shape carboxylic acid rich protein with a molecular weight of about 66000 Da. BSA has 584 amino acids and it contains three homologous domains which are connected by disulphide bonds [30, 31]. BSA adsorb on the hydrophobic surface of PMDS by combination of hydrophobic and electrostatic interactions between BSA molecules toward PDMS surface (Figure *8*,a)[30, 32]. Surface modification with BSA coating is an effective method to decrease cell adhesion to surface due to the presence of the anti-fouling layer of BSA on the surface [33].

**Figure 8.**
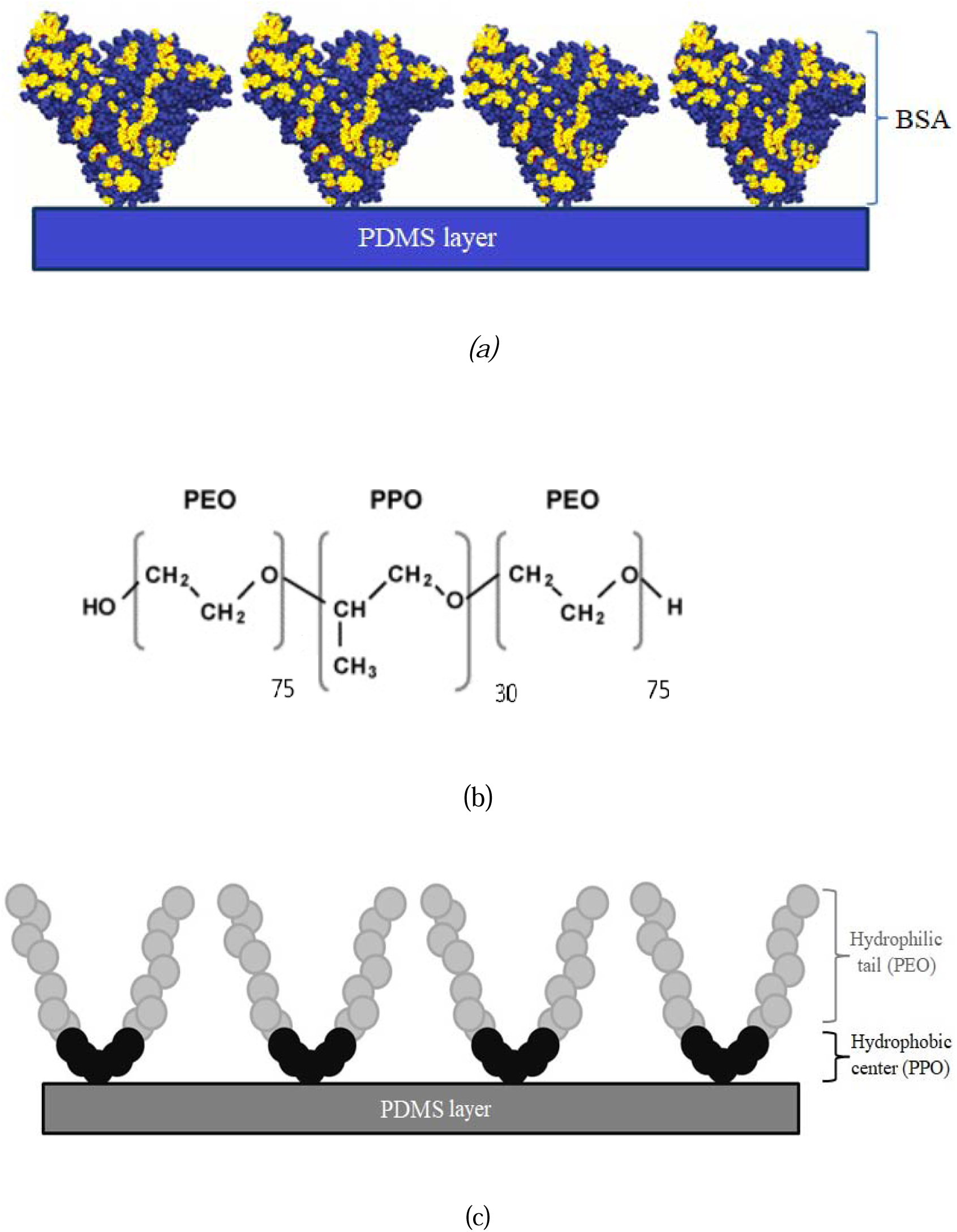
(a) Schematic of BSA molecules attaching and coating the PDMS surface (b) Chemical structure of Pluronic F-68, (c) Schematic of Pluronic F-68 molecules attaching and coating the PDMS surface

On the other hand, Pluronic F-68 ((PEO)75-(PPO)30-(PEO)75) (Figure 8, b) with the average molecular weight of 8400 Da was chosen from the family of triblock polymers [27]. It has been studied that Pluronic surfactants decrease protein adsorption and cell attachment to PMDS surface through spontaneous adsorption of their central hydrophobic poly (propylene oxide) (PPO) heads on the PDMS hydrophobic surface (Figure ***8***, c) which allows hydrophilic PEO tails extend out away from the surface [21, 27]

When the surface is treated with Pluronic F-68, the hydrophobic PPO head attach to the PDMS surface while the hydrophilic PEO chains extend freely into the interface and enhance surface hydrophilicity and make the surface resistant to protein/cell adhesion.

Since Pluronic is much more soluble in the water than that in the PDMS matrix, when PDMS contacts aqueous solutions of Pluronic, the hydrophilic PEO tails tend to migrate to the interface of PDMS and aqueous solution and the hydrophobic PPO heads adsorb onto hydrophobic PDMS surface via physisorption and hydrophobic interaction [27, 34]. Therefore, the hydrophobic PDMS surface will transform to a more hydrophilic and protein-resistant surface [21]. Schematic representation of Pluronic F-68 adsorbed to PDMS surface showed in Figure 3.

Figure 9 shows the shematic of the 2D view of a spheroid which have been used for calculations of the average mean diameter, spheroid size and the circularity of each spheroid. The average mean diameter is calculated as Dm= (Dmax+Dmin)/2 [26]. The spheroid’s sizes were presented as mean diameter ± standard deviation. The circularity of the spheroids has been reported by the Circularity=Dmin/Dmax.

**Figure 9.**
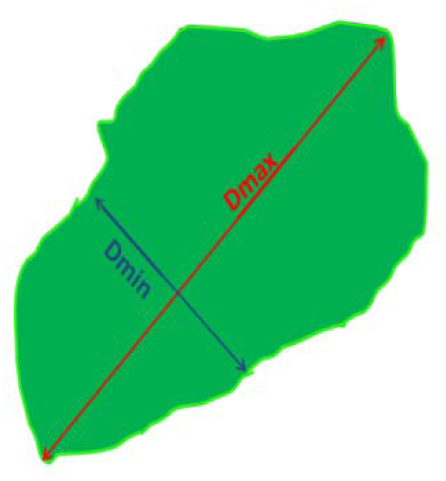
Schematic of the 2D view of the spheroid and the D-maximum and D-minimum which have been used in image J calculations to access the growth pattern and spheroid size over the 7 days periods

Figure 10 demonstrates the microscopic image of the microfluidic channels and chambers, a picture of the resin mold which is fabricated by stereolithography technic and a picture of the PDMS biochip which is used in this study.

**Figure 10.**
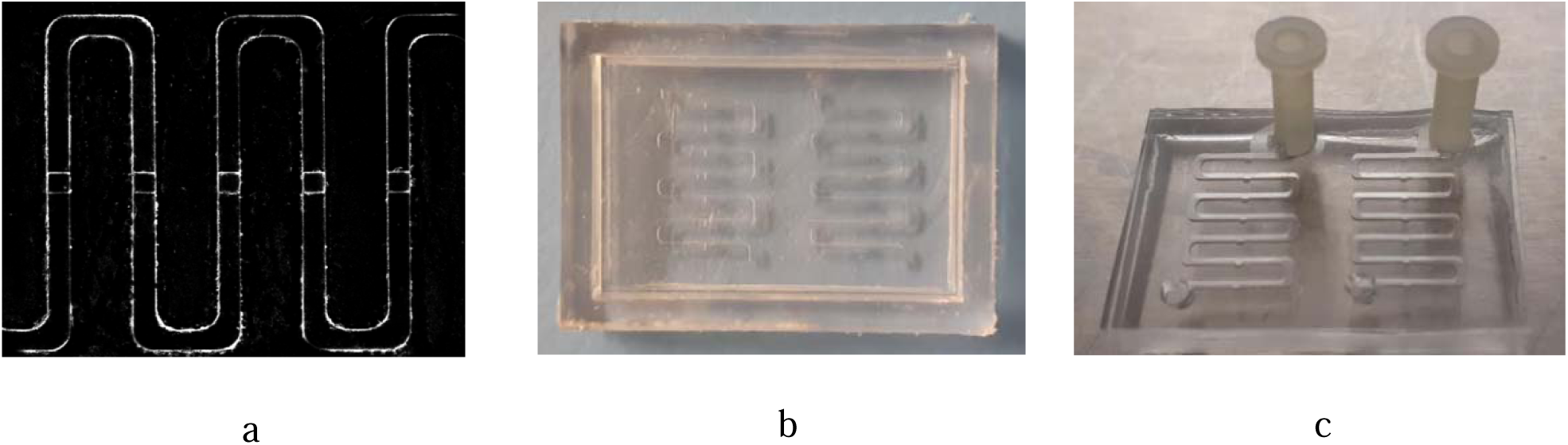
(a) Microscopic image of channels and chambers, (b) Picture of the 3D printed resin mold, (c) Picture of the PDMS microfluidic device with inlets

Figure 11 demonstrate the initial cell seeding process on biochip. Cell mixture is introduce to the inlet and some of the cells will trapped inside each well and the rest of the cells will exit through the outlet of the channel. As the inner surface of the PDMS biochip is treated with chemical coatings and render cell repellent properties, cells which are trapped inside the wells through sedimentation and containment will agglomerate together and proliferate to make 3D tumour spheroids on the day after cell seeding.

**Figure 11.**
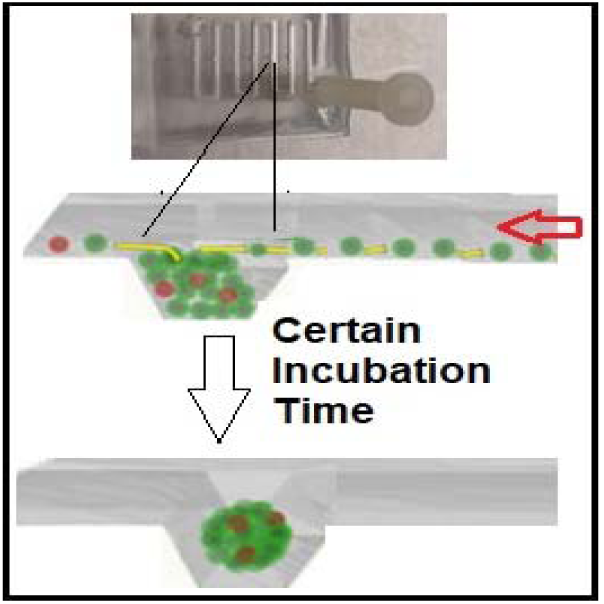
Schematic illustrations of the cell seeding process and 3D spheroid formation through sedimentation and containment

Figure 12 shows a comparison between the different treated PDMS surfaces in terms of the number of the spheroids forms on biochip. As discussed above, between 6 studied surfaces in this work, BSA 10% show the best cell repellent properties by increasing the number of possible spheroid formed on biochip. BSA 3% showed the least cell repellent properties and the less number of spheroid formed on-chip.

**Figure 12.**
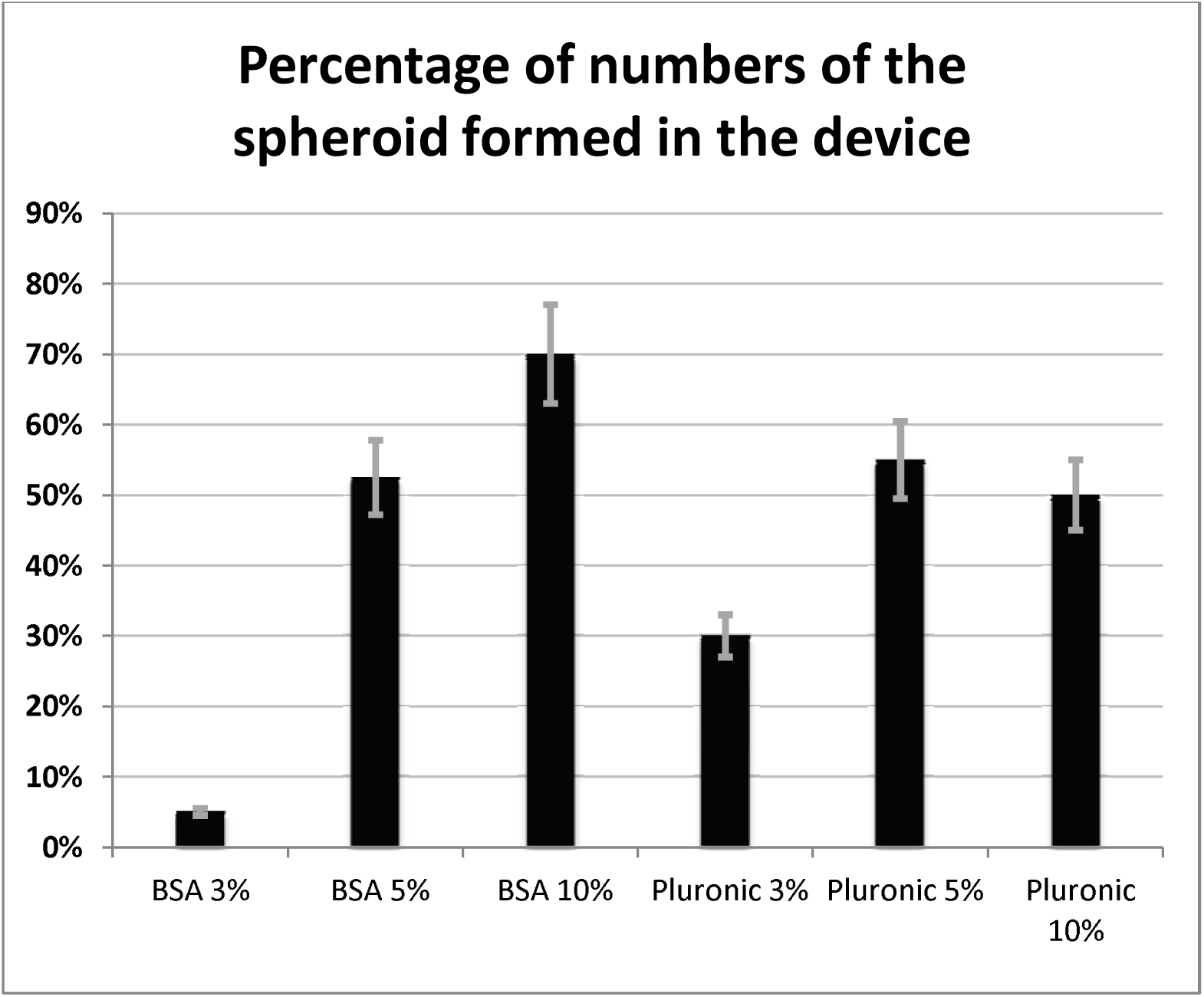
Percentage of the numbers of the spheroid formed in the device with different concentration of chemical coatings (BSA and Pluronic F-68.) (Error bars represent ± SE, n=3)

## Notes

### Competing Interest Statement

The authors have declared no competing interest.

